# Assesssing the role of humans in Greater Antillean land vertebrate extinctions: new insights from Cuba

**DOI:** 10.1101/2020.01.27.922237

**Authors:** Johanset Orihuela, Lázaro W. Viñola, Osvaldo Jiménez Vázquez, Alexis M. Mychajliw, Odlanyer Hernández de Lara, Logel Lorenzo, J. Angel Soto-Centeno

**Affiliations:** Department of Earth and Environment (Geosciences), Florida International University, Miami, Florida 33199, USA; Florida Museum of Natural History, University of Florida, Gainesville, FL 32611-7800; Gabinete de Arqueología de La Habana, Oficina del Historiador de La Habana, Cuba; Department of Rancho La Brea, La Brea Tar Pits & Museum, Los Angeles, CA 90036; Institute of Low Temperature Science, Hokkaido University, Sapporo, Japan 060-819; Laboratories of Molecular Anthropology and Microbiome Research, University of Oklahoma, Norman, OK 73019; Cuba Arqueológica, Progressus Heritage & Community Foundation, University of Florida, Gainesville, FL 32611; Fundación Antonio Núñez Jiménez de la Naturaleza y el Hombre, Jardines Bellamar, carretera las cuevas km 1½, Matanzas, Cuba; Department of Biological Sciences, Rutgers University, Newark, NJ 07102; Department of Mammalogy, American Museum of Natural History, New York, NY

**Keywords:** Anthropogenic factors, biodiversity, Caribbean, chronology, Cuba, extinction, extirpation, Holocene, island, vertebrates, West Indies

## Abstract

The Caribbean archipelago is a hotspot of biodiversity characterized by a high rate of extinction. Recent studies have examined these losses, but the causes of the Antillean Late Quaternary vertebrate extinctions, and especially the role of humans, are still unclear. Current results provide support for climate-related and human-induced extinctions, but often downplaying other complex bio-ecological factors that are difficult to model or to detect from the fossil and archaeological record. Here, we discuss Caribbean vertebrate extinctions and the potential role of humans derived from new and existing fossil and archaeological data from Cuba. Our results indicate that losses of Cuba’s native fauna occurred in three waves: one during the late Pleistocene and early Holocene, a second during the middle Holocene, and a third one during the last 2 ka, coinciding with the arrival of agroceramists and the early Europeans. The coexistence of now-extinct species with multiple cultural groups in Cuba for over 4 ka implies that Cuban indigenous non-ceramic cultures exerted far fewer extinction pressures to native fauna than the later agroceramists and Europeans that followed. This suggests a determinant value to increased technological sophistication and demographics as the most plausible effective extinction drivers.

## 1. Introduction

The Caribbean is a hotspot of biodiversity with some of the highest rates of extinction recorded throughout the Holocene (MacPhee and Flemming, 1999; Dávalos and Turvey, 2012). Studies on insular vertebrate biodiversity and their prehistoric remains have shown that the faunas present on this archipelago today are only a remnant of the richer species assemblages of the past (e.g. Morgan and Woods, 1986; Pregill et al., 1988). Recent biodiversity extinctions in the Caribbean are often attributed to natural phenomena (e.g. global climate change) and human effects, such as hunting, habitat alterations, and the introduction of exotic species (Morgan and Woods, 1986; Stoetzel et al., 2016; Borroto-Páez and Mancina, 2017). Globally, most island extinctions seem to coincide or follow the arrival of humans, although the magnitude and overlap of these events seem to vary depending on island size, human cultures, and the characteristics of the fauna (Turvey, 2009a, b; Steadman et al., 2015; Turvey et al., 2017). Other studies have further suggested that prehistoric and historic island extinctions likely represent a single continuous event linked to human activities (Lyons et al., 2016). Thus, as new fossil data becomes available, pinpointing the cause of these losses becomes more critical to further our understanding of the magnitude and tempo of species loss and examine clues to disentangle the causes of recent extinction. These efforts require fine-scale chronological evidence associated with detailed knowledge of human colonization events and local climatic changes.

The role of humans on Caribbean vertebrate extinctions, especially mammals and birds, has been debated for nearly four decades (e.g. Olson, 1982; Morgan and Woods, 1986; Pregill et al., 1994). While recent data has shed light upon the arguments of the cause, effect, and timing (Díaz-Franco 2004, 2011; Jiménez et al., 2005; Jull et al., 2004; Steadman et al., 2005; Silva et al., 2007; Orihuela, 2010; Orihuela and Tejedor, 2012; Soto-Centeno and Steadman, 2015; Cooke et al., 2017; Turvey et al., 2017; Borroto-Páez and Mancina, 2017), chronologic resolution and data consensus are still much needed from islands with the highest concentrations of biodiversity, such as Cuba.

Cuba, itself a sub-archipelago within the Greater Antilles, is the largest of all Caribbean islands and has one of the most diverse fossils and modern vertebrate records in the region (MacPhee and Fleming, 1999, MacPhee et al 1999b). Its high levels of endemism, geological complexity (Iturralde and MacPhee, 1999; Hedges, 2001), and complex history of multiple human colonization events (e.g. Napolitano et al., 2019) make its consideration crucial to understanding how Amerindian populations and their landscape modifications contributed to vertebrate extinctions (Silva et al., 2007). Nevertheless, Cuban zooarchaeological data has been rarely incorporated into current Caribbean biogeographical research programs and extinction syntheses. This has been exacerbated by the poor circulation and limited accessibility of relevant Cuban research or publications (e.g. Nuñez and Mayo, 1970; Pino and Castellanos, 1985; Díaz-Franco, 2004, 2011; Muñíz y Domínguez, 2014; Jiménez, 2015; Jiménez and Arrazcaeta, 2015). While prior studies have compiled LADs for 10 Cuban mammals, only three have direct dates (the eulipotyphlan *Nesophontes micrus*, and the sloths *Parocnus browni* and *Megalocnus rodens*; MacPhee et al., 1999, 2007; Turvey and Fritz, 2011), and associated radiocarbon or stratigraphic date estimates from Cuban zooarchaeological data has been scantly compiled. Additional illustrative examples from the Cuban literature that have been neglected include the well-documented cases of hutia “domestication” or captive husbandry, interisland exchange, or the introduction of exotic species by Amerindians (e.g. Pose et al., 1988; Jiménez and Milera, 2002; Díaz-Franco, 2004; Silva et al., 2007; Díaz-Franco and Jiménez, 2008).

Herein we provide novel details on Cuban vertebrate extinctions and highlight the potential role humans may have played in Cuban land vertebrate extinctions through the analysis of both new and existing radiometric and zoo-archeological data. Studies of Late Quaternary extinctions on both islands and continents typically evaluate the temporal overlap of faunal last appearance dates (LADs) representing the last time a species was detected in the fossil record, and the first time humans arrive, or first appearance date (FAD), representing the first dated sign of human activity (e.g. MacPhee and Flemming, 1999). We generated 17 new direct accelerator mass spectrometry AMS radiocarbon dates for endemic Cuban vertebrates (extinct and extant, terrestrial volant and non-volant), seven of which represent new direct LADs. Moreover, using detailed knowledge of the Cuban archaeological record and literature, we critically compare and contrast the timing of faunal extinctions and extirpation on the island and their relationship with the known Amerindian cultures that existed during those intervals.

## 2. Materials and methods

### 2.1 Regional setting

Cuba (22.025° N, −78.997° W; Fig. 1) is the largest island in the Caribbean, comprising an archipelago of about 4,000 islands and occupies about 46% of the total land area of the Caribbean (Woods and Sergile 2009). The island contains diverse habitats ranging from tropical dry to mesic forests, semideserts, and mountains that reach elevations no greater than 1974 m; conditions that contribute to the overall diversity species found there. We compiled fossil and archaeological data from 17 unique localities across western and central Cuba (Fig. 1, Table 1).

**Table 1.** Direct Accelerator Mass Spectrometer (AMS) radiocarbon (^14^C) dates for individual fossil remains from 17 unique localities across Cuba. All estimates represented as conventional ^14^C dates in chronological order. Intervals definitions: Late Amerindian (l Am), post Columbian (p Col), Amerindian (Am), Early Holocene (e HOL), Middle Holocene (m HOL), and Late Pleistocene (l PL).

| Taxa | Interval | Conventional<br>$^{14}\text{C}$ age (yr BP) | Calibrated $^{14}\text{C}$<br>age (cal BP, 2σ) | Sample No. | Site / Deposit | Source |
| --- | --- | --- | --- | --- | --- | --- |
| *Nesophontes micrus* | l Am–p Col | 490 ± 50 | 640–341 | Beta-115695 | Cueva Martin, Santi Spiritus (1) | MacPhee et al. 1999a |
| *Solenodon cubanus* | l Am | 650 ± 15 | 665–561 | UCIAMS-218808 | Cueva La Caja/Nesofontes, Mayabeque (2) | This paper |
| *Mesocapromys nanus* | l Am | 665 ± 20 | 670–562 | UCIAMS-218804 | Abra de Figueroa (owl roost), Matanzas (3) | This paper |
| *N. major* | l Am | 670 ± 15 | 670–565 | UCIAMS-218809 | Abra de Figueroa (owl roost), Matanzas (3) | This paper |
| *N. micrus* | l Am | 770 ± 50 | 790–653 | Beta-115696 | Solapa La Jaula (4) | MacPhee et al. 1999a |
| *S. cubanus* | l Am | 820 ± 15 | 760–690 | UCIAMS-218807 | Cueva Calero, Cantel, Matanzas (arch.) (5) | This paper |
| *N. micrus* | l Am | 840 ± 15 | 786–703 | UCIAMS-218810 | Abra de Figueroa (owl roost) (3) | This paper |
| *N. micrus* | l Am | 860 ± 30 | 901–695 | ICA-18B/1234 | Cueva El Gato Jíbaro, Matanzas (6) | This paper |
| *Antrozous koopmani* | l Am | 1150 ± 30 | 1174–979 | Beta-526491 | Cueva La Caja/Nesofontes, Mayabeque (2) | This paper |
| *Boromys torrei* | l Am | 1180 ± 15 | 1175–1061 | UCIAMS-218802 | Cueva La Caja/Nesofontes, Mayabeque (2) | This paper |
| *Homo sapiens* | l Am | 1190 ± 40 | 1255–984 | ICA-16B/0605 | Cueva del Muerto, Mayabeque (7) | Orihuela et al. 2016 |
| *A. koopmani* | l Am | 1280 ± 30 | 1288–1176 | Beta-526493 | Cueva La Caja/Nesofontes, Mayabeque (2) | This paper |
| *Artibeus anthonyi* | l Am | 1290 ± 30 | 1286–1180 | ICA 14B/1102 | Cueva La Caja/Nesofontes, Mayabeque (2) | Orihuela et al. unpub. |
| *A. koopmani* | l Am | 1300 ± 30 | 1291–1181 | Beta-526492 | Cueva La Caja/Nesofontes, Mayabeque (2) | This paper |
| *B. offella* | l Am | 1320 ± 15 | 1294–1187 | UCIAMS-218806 | Cuevas Blancas, Mayabeque (arch.) (8) | This paper |
| *Puslatrix arredondo* | l Am | 1390 ± 30 | 1348–1277 | ICA-16B/0606 | Cueva del Muerto, Mayabeque (7) | Jiménez et al. unpub. |
| *N. major* | l Am | 1418 ± 20 | 1348–1294 | Beta-392022 | Cueva La Caja/Nesofontes, Mayabeque (2) | Orihuela et al. unpub. |
| *H. sapiens* | l Am | 1570 ± 40 | 1548–1378 | ICA-19B/0508 | Cueva El Gato Jíbaro, Matanzas (6) | This paper |
| *Macrocapromys acevedo* | l Am | 1590 ± 30 | 1545–1409 | Beta-526495 | Cueva de Leonel, Trinidad (9) | This paper |
| *M. nanus* | l Am | 1675 ± 15 | 1613–1541 | UCIAMS-218803 | Cueva del Caracol (10) | This paper |
| *M. nanus* | l Am | 1775 ± 20 | 1780–1615 | UCIAMS-218805 | Cueva de los Paredones, Mayabeque (11) | This paper |
| *M. kraglievichi* | l Am | 1780 ± 50 | 1822–1569 | ICA-15B/0118 | Solapa del Megalocnus, Mayabeque (12) | This paper |
| *Phyllops vetus* | l Am | 1960 ± 30 | 1989–1830 | ICA 18B/0845 | Cueva La Caja/Nesofontes, Mayabeque (2) | Orihuela et al. unpub. |
| *Megalocnus rodens* | Am–m HOL | 4190 ± 40 | 4844–4584 | Beta-206173 | Solapa de Silex, Mayabeque (13) | MacPhee et al. 2007 |
| *Parocnus browni* | Am–m HOL | 4960 ± 280 | 6310–4975 | AA35290 | Las Breas de San Felipe, Matanzas (14) | Jull et al. 2004 |
| *A. anthonyi* | Am–m HOL | 5190 ± 40 | 6172–5773 | ICA-19B/0450 | Cueva El Gato Jíbaro, Matanzas (6) | This paper |
| *M. rodens* | Am–m HOL | 6250 ± 50 | 7270–7007 | B-115697 | Cueva Beruvides, Matanzas (15) | Steadman et al. 2005 |
| *P. browni* | l PL–e HOL | 10520 ± 440 | 13307–11138 | AA35291 | Las Breas de San Felipe, Matanzas (13) | Jull et al. 2004 |
| *P. browni* | l PL–e HOL | 11880 ± 420 | 15236–12973 | AA35292 | Las Breas de San Felipe, Matanzas (13) | Jull et al. 2004 |
| *Tyto noeli* | l PL | 17406 ± 160 | 21494–20590 | NA | El Abrón, Pinar del Río (16) | Suárez & Franco 2003 |
| *Pinus sp.* | l PL | >30000 ± 2000 | NA | L-130A | Casimbas de Ciego Montero, Cienfuegos (17) | Kulp et al. 1952 |
| *Unsp. Wood* | l PL | 25000 ± 2000 | 35503–25759 | L-130B | Casimbas de Ciego Montero, Cienfuegos (17) | Kulp et al. 1952 |

**Figure 1.**
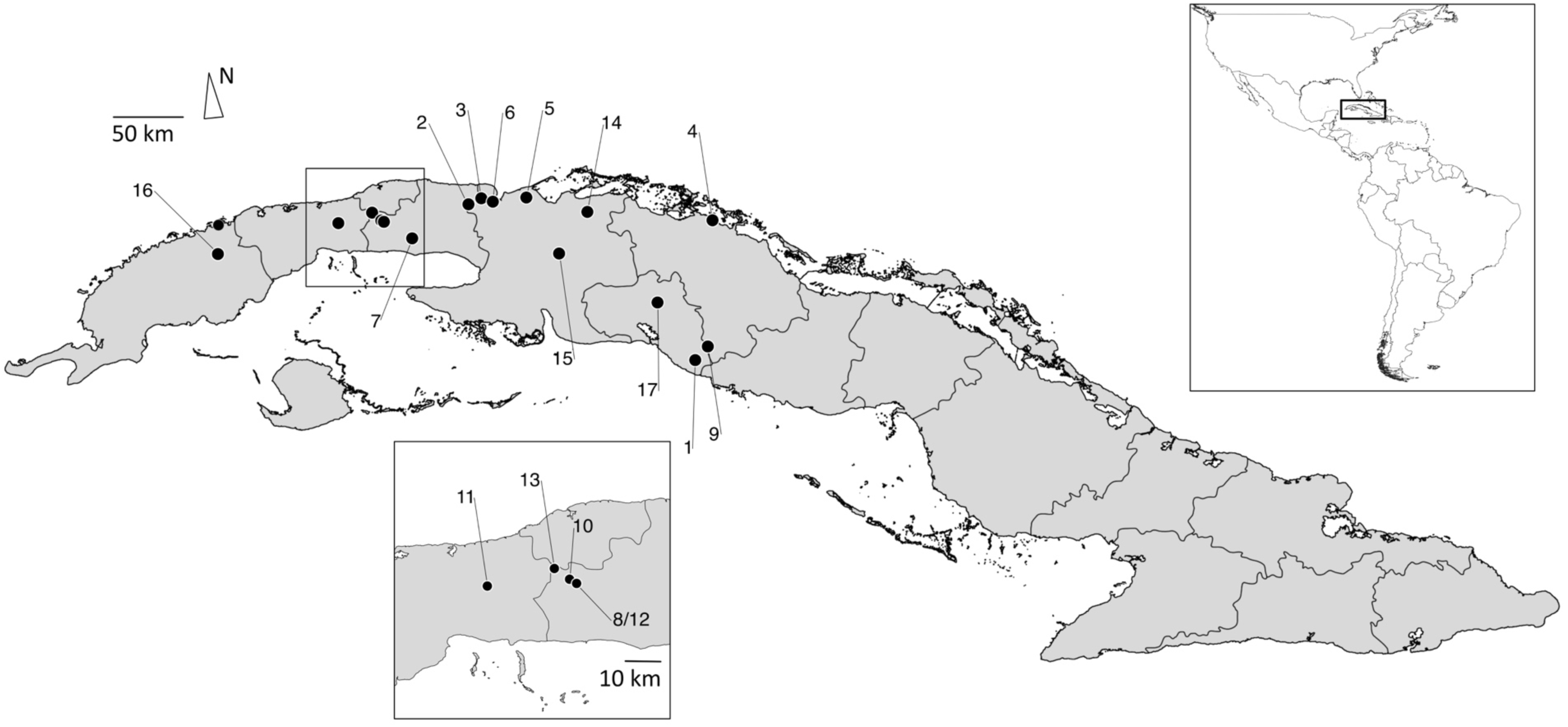
Map of Cuba highlighting the 17 unique fossil localities described in this study. Numbered labels represent locality names as listed in parenthesis in Table 1 and described in Supplementary Text S2 study site details. Cuba inset shows specific localities within Artemisa, La Habana, and Mayabeque provinces. Marker 8/12 in inset represents two nearby unique localities.

### 2.2 AMS Radiocarbon (^14^C) dated material

We present a dataset comprising seven direct last appearance ^14^C dates (LADs), cross-referenced with an additional 80 indirect chronologic taxa associations for extinct Cuban birds, mammals, and a tortoise extracted from available published sources; including previously unreported fauna from sites with known associated chronologies (Table 1 and Table S1). The new directly dated material included here was collected from several fossil-rich localities and complemented with material from 10 direct and 40 indirect records from the literature from (Table 1, Table S1, for study site details see Supplementary Text S1). Material excavated from these sites is deposited at the Museo Nacional de Historia Natural (MNHNCu) and at the Gabinete de Arqueología de la Oficina del Historiador (GAOHCu), both located in La Habana, Cuba. Accelerated Mass Spectrometry (AMS ^14^C) dating was run on purified bone collagen and performed at Beta Analytic (Miami, FL), International Chemical Analysis Inc. (ICA, Miami, FL), and the UC Irvine Keck Accelerator Mass Spectrometry facility (UCIAMS). Methods used at UCIAMS followed a modified Longin collagen extraction protocol (Brown et al., 1988) on specimens with suitable collagen yields and C: N ratios and were then ultrafiltered (> 30 kDa) to remove contaminants. All radiocarbon dates were calibrated using OxCal v. 4.3 (Bronk Ramsey, 2009) and the IntCal13 calibration curve (Reimer et al., 2013), as no marine samples were included. We report the median calibrated age (Cal BP) and the 95.4% range.

### 2.3 Taxonomy

For bats, we follow the systematic taxonomy of Silva (1974, 1976) and Balseiro (2011), except in considering *Mormoops magna* a valid extinct species. *Desmodus puntajudensis* is considered a synonym of *Desmodus rotundus* following Orihuela (2011). For terrestrial mammals, we follow Silva et al. (2007) and González et al. (2012) but consider the validity of two species of *Nesophontes* shrews (Condis et al., 2005; Rzebik-Kowalska and Woloszyn, 2012), and accept the new inclusion of the black-tailed hutia (*M. melanurus*) within the genus *Mesocapromys* (Upham and Borroto-Páez, 2017). We follow Orihuela (2019) for birds and Arana (2019) for Squamata. So far, *Chelonoidis cubensis* is the only extinct reptile registered from Cuba (Albury et al., 2018).

*Abbreviations*: AD, anno domini or common era. BC, before Christ or before the common era. BP, before the present (datum AD 1950). Cal., calibrated age. Cal BP, median radiocarbon calibrated age before the present. EED, effective extinction date. eHOL, early Holocene. Ka, kilo-years. LAD, last appearance date. LGM, Last Glacial Maximum. lHOL, late Holocene. PHT, Pleistocene-Holocene transition. Rdcy, radiocarbon years before the present.

### 2.4 Chronologic intervals and last appearance dates (LAD)

For ease of communication and consistency with previous literature, we used chronologic intervals names in Soto-Centeno and Steadman (2015). The first interval is the Last Glacial Maximum or Late Pleistocene (ca. 21–11.7 ka), which culminated with the Pleistocene-Holocene transition (PHT, ca. 11.7 ka; Curtis et al., 2001). The following intervals are the early Holocene (eHOL, ca. 11.7–8.2 ka), the middle Holocene (mHOL, ca. 8.2–4.2 ka), and the late Holocene (lHOL, < 4.2 ka). We also separately demarcate the Amerindian interval, defined as the beginning of the transition between the middle to late Holocene (mHOL to lHOL, < ∼6 ka), and spans the entire late Holocene. A final and fifth stage is the post-Columbian interval beginning in the last 500 BP (AD 1492–1500 to the present), interchangeably used in the literature as the European or Historic interval (e.g. Morgan and Woods, 1986; Soto-Centeno and Steadman, 2015; Cooke et al., 2017).

#### 2.4.1 A rationale for a subdivision of the Amerindian interval

As humans first arrived in Cuba after the middle Holocene (mHOL, < ∼6 ka; Cooper, 2010; Napolitano et al., 2019), we subdivide the Amerindian interval into two subintervals to better discuss LADs in the context of different cultural practices and degrees of environmental alteration, as they relate to patterns of extinction and extirpation. These two intervals are the early Amerindian (∼6–1.5 ka BP) and the late Amerindian (1500–500 BP). Our use of this specific scheme focuses attention on the cultural groups that could have importantly affected the natural environments of the island at a given chronological interval and provides a better resolution to the timing of extinction or extirpation. An expanded rationale for these subintervals, including the difference between “archaic” and “ceramic” cultures, climate-related information, and a description of the sites and deposits are provided in the Supplementary Text S2.

## 3. Results

The direct dates we report represent 27% (16 of 59) of the known extinct land vertebrate fauna, including four extant but critically endangered mammal species. Within these 59 vertebrates are 38 birds and one reptile, which together comprise the record of extinct and extirpated land vertebrate species so far registered from the Quaternary of Cuba. We provide direct and indirect LADs for 95% (20 of 21) of the known Cuban extinct land mammal fauna, including bats. This overall mammalian fauna is composed of two echimyid rodents, four capromyid rodents, four sloths, three eulipotyphlans, one monkey and seven bats (Fig. 2–4, Table 1). Moreover, we provide new direct radiocarbon dates on Amerindian human remains that contribute to a better understanding of the contexts of association for several species that lack direct dates (Fig. 2, Table S1), and provide further support to formerly estimated LADs (e.g. for the extirpated bat *Mormoops megalophylla* from Cueva del Gato Jíbaro in Orihuela and Tejedor, 2012; see Table 1, Table S1).

**Figure 2.**
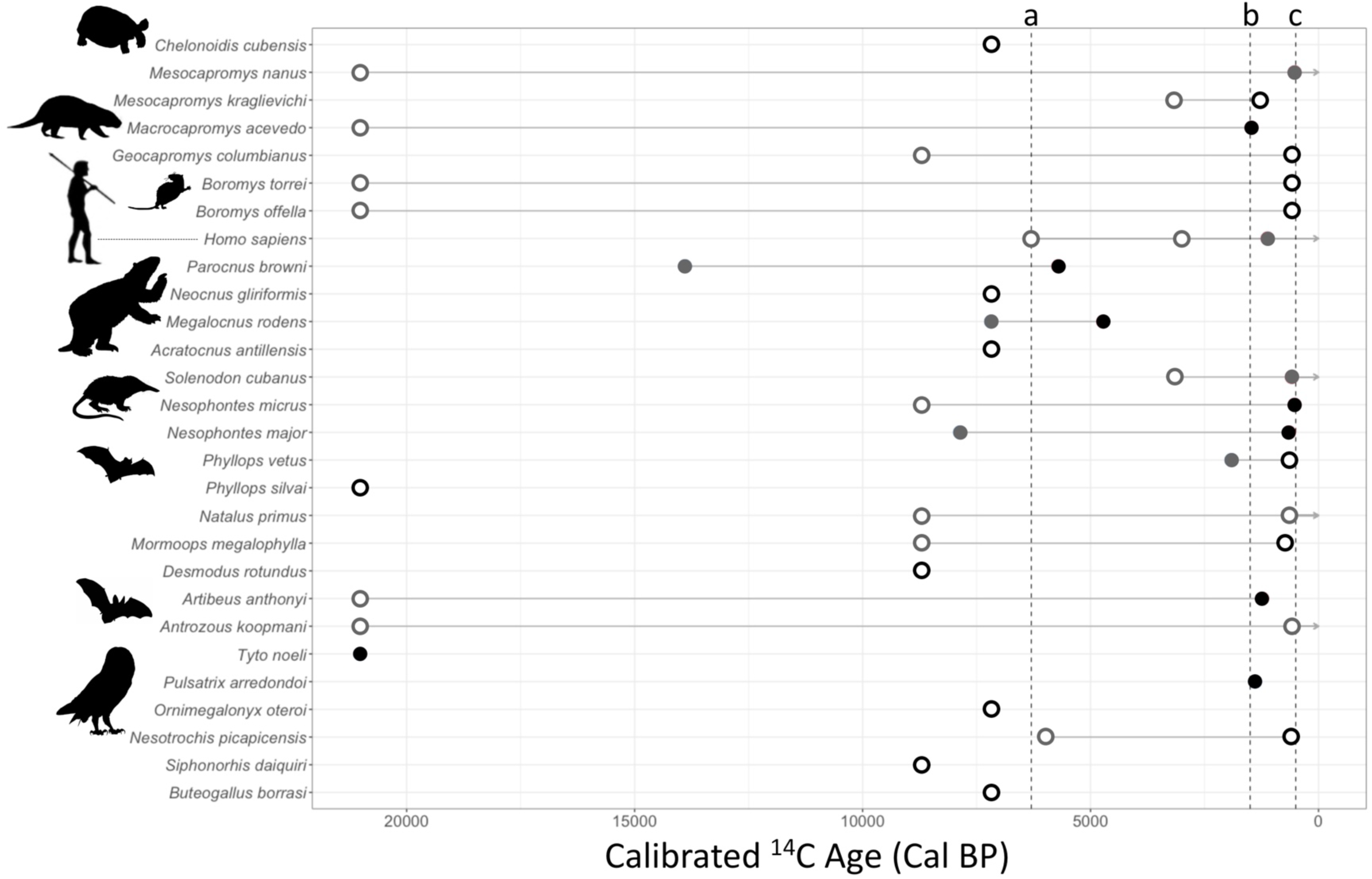
Chronology of extinction of Cuban vertebrates based on calibrated AMS radiocarbon (^14^C) dates. Taxonomic groups are indicated by a silhouette (respectively: Testudines, Rodentia, Primates, Pilosa, Eulipotiphla, Chiroptera, Aves). Circles represent radiocarbon dates: closed circle = direct date, open circle = indirect date, gray = earliest appearance in the fossil record, black = last appearance date (LAD). Horizontal bars are used to highlight the existence of a taxon over time and arrows indicate extant taxa. Vertical dashed lines show the Early Amerindian (a = ∼6ky), Late Amerindian (b = ∼1500y), and post-Columbian (c = <500y) subintervals (see Methods and Supplementary Text S2).

**Figure 3.**
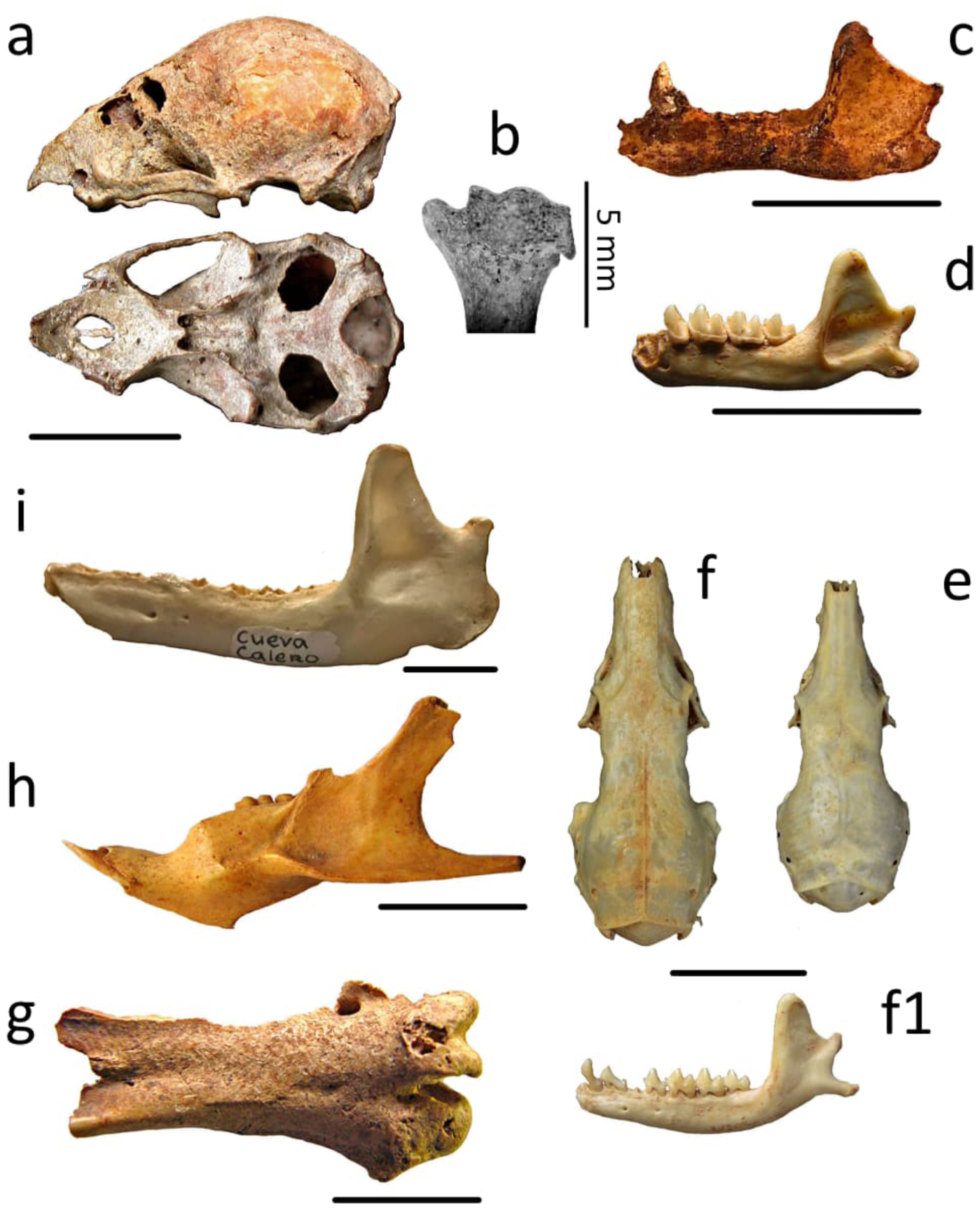
Representative extinct and endangered vertebrate fauna from Cuba associated with radiocarbon LADs. Clockwise from top left: a = *Desmodus rotundus* skull (Post-Columbian, Cueva de los Nesofontes); b = *Desmodus rotundus* distal humerus (Post-Columbian, Cueva de los Nesofontes; an attempt to directly date this specimen was unsuccessful); c = *Artibeus anthonyi* left hemimandible (paleontological context, ∼5190 BP, Cueva del Gato Jíbaro); d = *Antrozous koopmani* left hemimandible (∼1150 BP, Cueva de los Nesofontes); e = *Nesophontes micrus* skull (late Amerindian, Cueva de los Nesofontes); f and f1 = *Nesophontes major* skull and hemimandible (∼1418 BP, Cueva de los Nesofontes); g = *Pulsatrix arredondoi* tarsus (∼1390 BP, Cueva del Muerto); h = *Mesocapromys kraglievichi* left hemimandible (late Amerindian, Cueva de los Nesofontes); i = *Solenodon cubanus*, left hemimandible (∼820 BP, Cueva de Calero). All bars scaled to 10 mm, except in b, which is as indicated.

**Figure 4.**
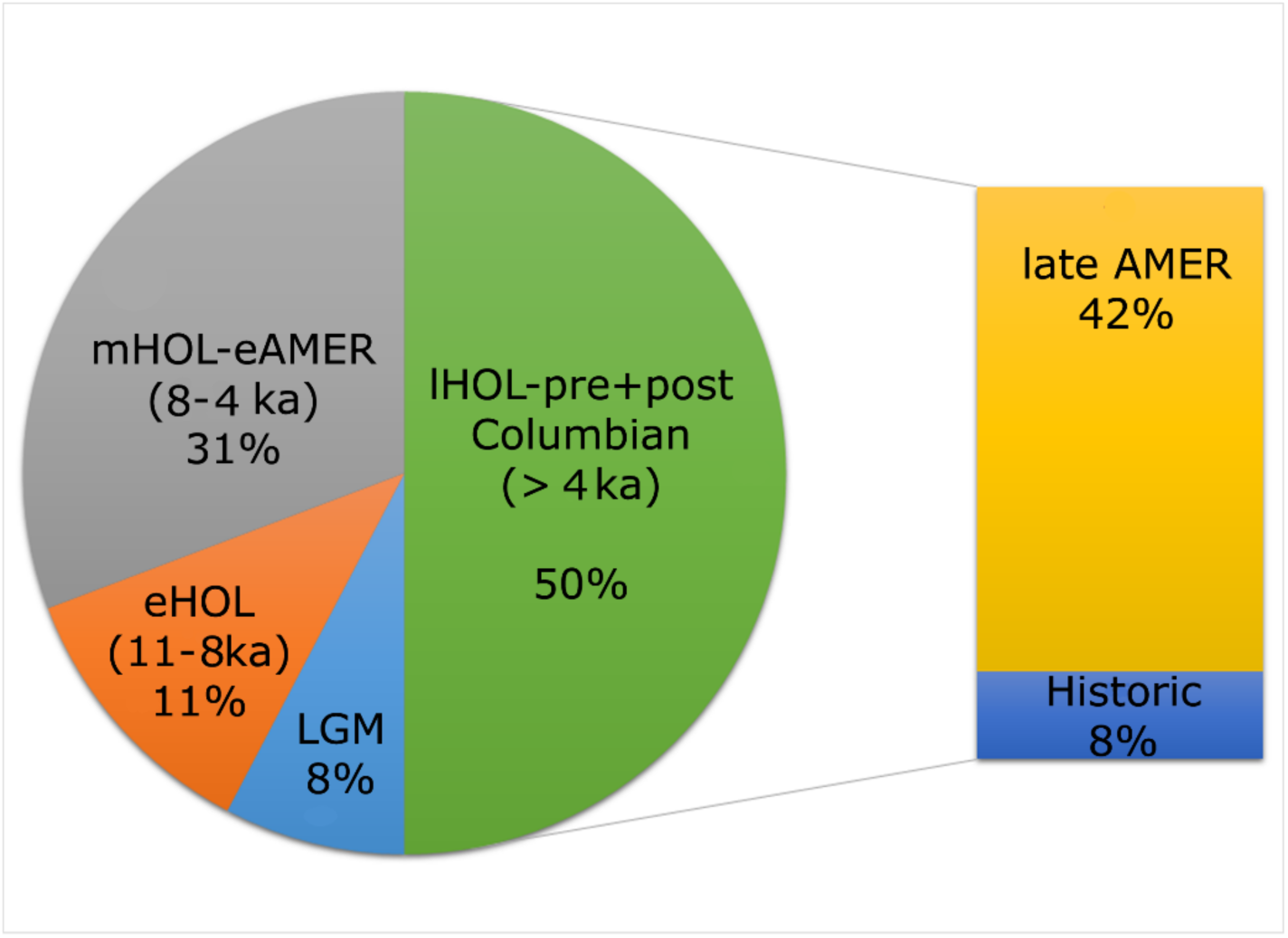
Proportion of extinctinct taxa per anthropogenic interval in Cuba. Half of the documented LADs postdate 4ka and about 42% of these date within the later Amerindian interval defined here (see Methods and Supplementary Text S2).

Of the chronologically analyzed extinct and extirpated vertebrates, 44.4% (12 of 27) dated well into the late Amerindian (1500–500 BP; Fig. 2, 4, Table 1, Table S1–S2). Only two (7.4%) of the 27 currently registered extinct species have LADs within the Late Pleistocene (ca. 21–11.7 ka), 11.1% (3 of 27) occur during the early Holocene (eHOL, ca. 11.7–8.2 ka), 33.3% (9 of 27) occur during the middle Holocene (mHOL, ca. 8.2–4.2 ka), and 48.1% (13 of 27) occur during the Late Holocene (lHOL, ca. > 4.2 ka; Orihuela, 2019). When examining these 13 late Holocene species based on Amerindian subintervals, only three species (*Macrocapromys acevedo, Parocnus browni*, and *Megalocnus rodens*) had LADs on the early Amerindian subinterval (ca. 6–1.5ka), whereas 10 dated within the late Amerindian (ca. 1.5ka–500 BP). At least four of these 13 species could have had EEDs in the post-Columbian interval (ca. 500 BP– present): *Boromys torrei, B. offella, Geocapromys columbianus* and *Nesotrochis picapicensis* (Fig. 2, 4, Table 1; Tables S1–S3). So far, only 5.4% (2 of 38) currently known extinct and extirpated Cuban birds, have direct radiometric LADs; one at the LGM and one in the late Amerindian (Orihuela, 2019). There are no known extinct amphibians described so far, and only one reptile, *Chelonoidis cubensis* (Albury et al., 2018).

For mammals, we provide direct dates from specimens of the extinct eulipotiphlan insectivores *Nesophontes major* and *Nesophontes micrus*, the extinct fruit bats *Artibeus anthonyi* and *Phyllops vetus*, the likely extinct *Antrozous koopmani*, plus the pygmy hutia *Mesocarpomys nanus* and *Solenodon cubanus*, the extinct rodents *Boromys torrei, B. offella, Macrocapromys acevedo*, and *Mesocapromys kraglievichi*. We also provide associated age estimates for the extirpated bats *Mormoops megalophylla* and *Desmodus rotundus*, and the hutia *Geocapromys columbianus* based on new indirect radiocarbon dates (Table S1). All their LADs are also well within the late Amerindian subinterval, supporting formerly presumed extinction estimates within the Late Holocene and confirming the coexistence of native fauna with humans after their initial arrival (Jiménez et al., 2005; Orihuela, 2010; Orihuela and Tejedor, 2012; Orihuela et al., 2020). New associated LADs are also provided for the sloths *Acratocnus antillensis* and *Neocnus gliriformis*, and the extinct tortoise *Chelonoidis cubensis* based on associated date estimates, support their survival up to the early Amerindian, middle Holocene (Table 1, Table S1).

Remaining uncertainties still exist in the Cuban quaternary fossil record. Two extinct bats (*Cubanycteris silvai* and *Pteronotus pristinus*) do not have direct or indirect radiocarbon dates available to establish LAD estimates. Their extinction interval is assumed to be Late Pleistocene based on the interpretation of the authors (Silva 1974, 1979; Mancina and García-Rivera, 2005; Suárez and Díaz-Franco, 2003), but likely that these could be extended at least to within the early Holocene (eHOL, ca. 11.7–8.2 ka), based on fauna association and other data discussed in Condis (2005) and Fiol (2015). A data gap also limits the constraint of the extinction of the monkey *Paraloutta varonai* and the eulipotyphlan insectivore *Solenodon arredondoi* (Salgado et al., 1990; MacPhee et al., 1996a; Morgan and Ottenwalder, 1993; Jiménez, 2015), but an early Holocene extinction is also plausible for these taxa based on their fauna associations.

Generally, eulipotyphlan insectivores and native rodents yielded LADs that are borderline to the post-Columbian interval, which seem to corroborate former assumptions of their extinction during the historic interval (Guarch, 1982, 1984; Pino, 2012; Fig. 2, Table 1, Table S1–S2). Direct and indirect dates for extant but critically endangered species, such as *Mesocapromys nanus, Solenodon cubanus*, and the bats *Antrozous koopmani* and *Natalus primus*, suggest that these species had a much wider distribution throughout western Cuba up to the time of European colonization. In this sense, these data represent ecological baselines that can be used to evaluate the trajectory of range collapse or earlier stages of extinction processes.

## 4. Discussion

Extinctions have had a significant role in shaping the living vertebrate diversity of islands globally, including the Caribbean and the Cuban sub-archipelago within it. Detailed radiochronologies and robust estimates of last appearance dates (LADs) of large fossil assemblages allow us to analyze changes in community composition and the timing of community turnover events in association with the anthropogenic perturbations that drove them. Previous studies debating the cause of extinction of Cuban vertebrates have typically done so despite lacking appropriate resolution of chronological data. For the first time in this system, we assessed extinction events using sufficient radiocarbon estimates of 56% of all extinct taxa documented for the island. We found that nearly half of Cuban species within our study went extinct during an interval of active human presence (lHOL, ca. > 4.2 ka). Our results reveal that at least five species disappeared before the arrival of humans, whereas potentially 7–8 species went extinct during the middle Holocene following human arrival after ∼6 ka. Most of the extinct species in Table 1 survived the Pleistocene-Holocene transition (PHT, ca. 11.7 ka) and several thousand years of human habitation of Cuba (Fig. 2, Table S3).

Up to 90% (22 of 27) of the vertebrate fauna we studied have direct and indirect LADs younger than eHOL (ca. 8.2 ky; Fig. 2, Table 1, Table S2–S3). The majority of these (66.6%) have LADs well into the late Amerindian and post-Columbian intervals, coinciding to a time near or after the arrival of the agroceramist culture groups in Cuba (ca. ∼ 1500 BP). These data support the hypothesis that lost species persisted for thousands of years after the onset of warming climate conditions of the Holocene. Furthermore, our results show persistence for over 4 ky of Amerindian presence in Cuba, which is congruent with estimates from previous studies on other islands and also supports previous late Holocene extinction estimates in Cuba (MacPhee et al. 1999; Díaz-Franco 2004, 2011; Jiménez et al., 2005; Orihuela, 2010; Orihuela and Tejedor, 2012; Soto-Centeno and Steadman, 2015; Borroto-Páez and Mancina, 2017).

The earliest Amerindians to arrive in Cuba were the so-called “archaic” pre-Arawak that reached the island during the mid-Holocene (Cooper, 2010; Cooper and Thomas, 2011; Roksandic et al., 2015; Ulloa and Valcárcel, 2016, 2019). These earliest indigenous colonists were preceramic hunter-gatherer groups with estimated low demographics and they exploited a wide range of ecosystems within the archipelago (Guarch et al., 1995; González Herrera, 2008, 2018; Ulloa and Valcárcel, 2019).

Early Amerindian culture middens show a preference for coastal and riverine resources, predominantly mollusks and fish (Pino, 1978, 2012), with no direct evidence of consumption of large mammals such as sloths and rodents. Rodents were secondarily exploited, especially the larger hutias such as *Capromys pilorides* and *Geocapromys columbianus*, and the medium to smaller sized *Mysateles prehensilis, Mesocapromys melanurus, Boromys* spp. (Kozlowski, 1974; Pino, 1978, 2012; Córdova-Medina et al., 1997; Reyes, 1997; Jiménez and Arrazcaeta, 2015), whereas the eulipotyphlans *Solenodon* and *Nesophontes* were probably occasional supplemental food items (Martínez Arango, 1968; Pino, 1978, 2012; Martínez et al., 1993).

Although these preceramic indigenous groups were dietarily, ecologically and culturally diverse, it is unlikely that they had a major impact on Cuban fauna due to their incipient technological sophistication and demographics. Many of the larger mammals (e.g, sloths, monkeys) and the land tortoise (*Chelonoidis cubensis*) seem to have disappeared during the mHOL, near the time of first human arrival. Despite our increased temporal resolution, it is still difficult to discern how early colonists could have perturbed these vertebrate communities. For cavernicolous bats, for example, frequent human visits to caves likely could have affected their colonies (Silva, 1979; Mancina et al., 2007). The level of disturbance on bats could have been further exacerbated if cave fires are considered because smoke can lead to high mortality or complete extirpation. Nonetheless, direct evidence for these mass deaths or disturbance is lacking from the current record in Cuba (Orihuela and Tejedor, 2012).

Sloth remains are rare from midden deposits attributed to early Amerindian groups. While the context and age of the evidence are still debated, sloths have been reported from at least a dozen archaeological sites (Pino y Castellanos, 1985; Jiménez and Arredondo, 2011:208; Díaz-Franco, 2004, 2011; Pino, 2012; Arredondo and Villavicencio, 2011). Taphonomic evidence of human predation in the form of tool and cut marks has been scant and remains questionable (Arredondo and Villavicencio, 2011; Jiménez and Arredondo, 2011; Orihuela et al., 2016). LADs for *Megalocnus rodens* and *Parocnus browni* and several of these other species (Table 1, Table S1) suggest that they coexisted in time but there is yet no conclusive evidence that sloths were predated by early Amerindians (Jull et al. 2004; MacPhee et al. 2007).

Other groups of hunter-fisher-gatherer Amerindians are also present in the Cuban archaeological record with radiocarbon dates between 2500 BC and AD ∼1500 (Guarch, 1990; Guarch et al., 1995; Cooper, 2007, 2010; Chinique and Rodriguez, 2012; Roksandic et al., 2015; Chinique et al., 2015; Chinique et al., 2019; see Supplementary Text S2). At least in Cuba, these Amerindians had a varied toolkit and likely practiced incipient agriculture (i.e. use of cultigens and wild varieties; see Chinique et al. 2015, 2016, 2019) with comparable adaptations but with slightly larger demographics given that some groups coexisted with the Taíno and Europeans that arrived later around 1500–500 BP, respectively (Fernández Oviedo, 1535; Las Casas, 1875; Guarch,1978, 1982; Dacal, 1980; Rouse, 1992; Torres, 2006). The hunter-fisher-gatherer midden remains, and direct isotope analyses indicate that they had variable diets acquired in diverse habitats (Chinique and Rodriguez, 2012; Chinique et al., 2015, 2016, 2019). Some groups were dependent on land mammals, in addition to marine/estuarine organisms (op. cit.). Rodents, particularly *Capromys pilorides, G. columbianus*, and the smaller spiny rats *Boromys* spp., were an important component of this culture’s diet (Guarch, 1982; Córdova-Medina, 1993; Reyes, 1997; Díaz-Franco, 2004; Jiménez and Arredondo, 2011; Pino, 2012; Colten and Worthington, 2018). It is likely these rodents survived past the post-Columbian interval (> AD 1500) even though most have LADs within the late Amerindian subinterval (Fig. 2).

The first agroceramists collectively called Taíno arrived in Cuba ∼1500 BP (AD 800– 900; Guarch, 1978; Rouse, 1992; Guarch et al., 1995; Valcárcel, 2002, 2008; Torres, 2006). Radiocarbon dates between the “archaic” pre-Arawak groups and Taíno groups overlap, supporting coexistence between different cultural groups at least in some areas of the archipelago as observed by early European chronicles (Fernández de Oviedo, 1535; Las Casas, 1875; Dacal, 1980; Chinique et al., 2016, 2019; Orihuela et al., 2017).

The Taíno hunted, fished, gathered, practiced land clearing for agriculture and lived in a wide range of sites that they could have fully exploited (Guarch, 1978; Rouse, 1992; Torres, 2006). They had a strong impact on the environment as evidenced by the introduction of domestic dogs (Arredondo, 1981; Wing and Scudder, 1983; Jiménez y Fernández-Milera, 2002; Newsom y Wing, 2004; Borroto-Páez, 2011; Hofman et al., 2011; Grouard et al., 2013; Laffoon et al., 2013). Although, it is not known when exactly the Taino introduced the domestic dog on the island. Chronicler accounts on *Canis lupus familiaris* indicate that it was already widely dispersed within the island by ∼1450–1530 AD when other domestic dogs were introduced by the Spanish colonists (Rodríguez-Durán y Santiago, 2014). Some taphonomic evidence supports canid predation on the native vertebrate fauna, including *Solenodon* and large rodents, and also feeding on human middens in both Taino and European contexts (Jiménez and Arrendondo, 2011; Orihuela et al., 2016).

Perhaps the most important interval for human-wildlife interactions in Cuba began during Taino’s coexistence with Europeans (< 1492 AD; Marrero, 1972; Morgan and Woods, 1986; MacPhee et al., 1999a/b; Valcárcel, 2012). These two waves of human colonization serially combined (i.e. agroceramists followed by Europeans) may have served as a series of continuous environmental perturbations through direct habitat destruction and the introduction of non-native species including house mice (*Mus musculus*), rats (*Rattus* spp.), domestic dogs (*C. lupus familiaris*), pigs (*Sus scrofa*), rabbits, New World monkeys, mongoose (*Herpestes auropunctatus*), and others (Borroto-Páez, 2011). These introduced species likely intensified pressure on native vertebrate populations like *S. cubanus, Boromys* spp., and *Nesophontes* spp. (Morgan and Ottenwalder, 1993; MacPhee et al., 1999; Silva et al., 2007; Jiménez and Arredondo, 2011).

Habitat loss, the introduction of non-native species, direct hunting, and anthropogenic climate change are some of the most important indirect anthropogenic drivers of modern extinctions in the Caribbean (IUCN, 2019). Many of these drivers have their roots in the Holocene, and we hypothesize that the combination of the latter agroceramist (Taino) and European colonization explains most species’ LADs (i.e. disappearances) during the last 2000 years (Steadman et al., 1984; Pregill et al., 1988; 1994; Turvey et al., 2007; Soto-Centeno and Steadman, 2015). Evidence that all the species that disappeared during the late Amerindian and post-Columbian intervals had survived over 4000 years of previous human inhabitation on the island, as in other areas of the Greater Antilles (e.g. Soto-Centeno and Steadman, 2015; Stoetzel et al., 2016, Soto-Centeno et al., 2017), validates the importance of the combination of threats brought by new colonizers with increased demographics, better environmental knowledge, and the technology to exploit it.

## 5. Conclusion

Our new radiocarbon dates and review support the hypothesis that extinctions in Cuba unfolded across multiple episodes during the Holocene. The majority of these extinction events are reflected in the direct and indirect LADs that coincide with the arrival of agroceramist groups and others later on, in the subinterval we defined as the late Amerindian (> 1.5 ka). Our data suggest that at least in Cuba, nearly half of the extinct land mammals (i.e. 44 %), and at least some avian extinctions, occurred during the last 1500 years. Making the late Amerindian and post-Columbian extinction episodes the most significant in Cuba’s late Quaternary. Historically, the first two centuries of colonialism in Cuba were not considered as having a strong impact on the fauna because large deforestation and environmental modifications began after the XVIII century.

Although a seemingly important extinction event is observed from LADs within the early Amerindian intervals (i.e. 33% of the 27 analyzed taxa have LADs between 6–1.5 ka), there is no evidence for a megafaunal overkill or blitzkrieg extinction in Cuba. It is possible that this apparent extinction episode is an artifact of our lack of data since taxa with LADs within this interval could have persisted later in time. Further, not all extinctions were driven by humans because some species did go extinct before human arrival to the archipelago (e.g. the bats *Pteronotus pristinus, Cubanycteris silvai, Phyllops silvai*, and the monkey *Paralouatta varonai*). A cascade effect that, acted concomitantly between climatic and anthropogenic factors, could have resulted in the asynchronous extinction of native land mammal and avian fauna extinctions we see that took place during the middle Holocene.

The LADs discussed herein indicate that nearly half of the species we sampled and listed survived over 4000 years of human coexistence in Cuba. The earliest human communities likely affected some groups (e.g. large terrestrial birds and sloths), either directly or indirectly, but caused little if any major environmental alterations in Cuba. Half of the species more frequently exploited and consumed survived. This condition dramatically changed after the arrival of the first agroceramist (Taíno) groups and was amplified later by the European colonists. It seems apparent that agroceramist and European colonizers could have played a decisive role in the direct or indirect extinction, extirpation or endangerment of Cuba’s native fauna. The effect of climate-related factors cannot be decoupled from anthropogenic ones precisely because they are concomitant from the onset of any human colonization, and do not disappear thereafter. Before human arrival, climate and bio-ecological-related factors likely played the most important role in determining geographical distribution, local or final extinction, and this explains extinctions during the Late Pleistocene to Holocene transition (see Supplementary discussions). This changed significantly at the arrival of Amerindians and later European colonists. Then, climate-related factors likely became secondary or less of a driver of changes in distribution or extinction in comparison to the more drastic human-related threats that are still present.

The consideration of the complex combination of climate and anthropogenic factors combined can provide a more realistic explanation of the extinction episodes observed after the Amerindian intervals. While some species may have survived previous changes in climate by shifting their distributions, they finally came to extirpation or extinction due to other causes, many of which remain unexplored (Supplementary Text S2). Based on the information at hand, the cause for Cuban vertebrate mid to late Holocene extinctions cannot be solely attributed to either climate or human-related drivers, either direct or indirect impacts, but more likely resulted from a combination of complex factors, including variables that we cannot model for, or that are not yet detected in the fossil record. Understanding the timing and the complexity of the causes leading to the extirpation or global extinction of native fauna is crucial to develop future protection and conservation programs, which must consider human demographic and technological growth, deforestation, climate change, and the individual species responses to these pressures.

## Acknowledgments

JO thanks Sigma Xi for providing funding for some of the analyses (grant id: G20141015720649). We thank Leonel Perez Orozco, Candido Santana, and Ricardo Viera for field assistance, discussions and logistics. Ercilio Vento, Stephen Díaz Franco and Abel Hernández for providing specimens used in this study. The *Consejo Nacional de Patrimonio Cultural, Ministerio de Cultura de Cuba* for excavation/exportation permits, and Jorge Garcell and Jaime Triana for all their help in securing them. We thank Adrian Tejedor, Herman Benitez, Nick Czaplewski, Gary Morgan, Jorge Ulloa, Roberto Valcárcel, Mathew Peros, Jago Cooper, Yadira Chinique, Mirjana Roskandic, Bronislaw Woloszyn and Ross MacPhee for literature and guidance. JASC thanks R.D. Barrilito for support. Radiocarbon work was partly funded by a Rutgers University Research Council Award to JASC. AMM was supported by a Japan Society for the Promotion of Science Short-Term Postdoctoral Fellowship. In figure 1, images for rat, sloth, solenodon, and vespertilionid bat were obtained from phylopic.org and used with permission under a creative commons license; other images produced by JASC.

## Notes

#### Summary of Updates

Correction on Figure 1.

